# Design of DNA Origami Diamond Photonic Crystals

**DOI:** 10.1101/2019.12.17.880302

**Authors:** Sung Hun Park, Haedong Park, Kahyun Hur, Seungwoo Lee

**Affiliations:** KU-KIST Graduate School of Converging Science and Technology, Korea University, Seoul 02841, Republic of Korea; Materials and Life Science Research Division, Korea Institute of Science and Technology, Seoul 02792, Republic of Korea; Department of Biomicrosystem Technology, Korea University, Seoul 02841, Republic of Korea; KU Photonics Center, Korea University, Seoul 02841, Republic of Korea

**Keywords:** DNA origami, Photonic crystals, Diamond lattice, Photonic bandgap (PBG), Effective elastic moduli

## Abstract

Self-assembled photonic crystals have proven to be a fascinating class of photonic materials for non-absorbing structural colorizations over large areas and in diverse relevant applications, including tools for on-chip spectrometers and biosensors, platforms for reflective displays, and templates for energy devices. The most prevalent building blocks for the self-assembly of photonic crystals are spherical colloids and block copolymers (BCPs) due to the generic appeal of these materials, which can be crafted into large-area 3D lattices. However, due to the intrinsic limitations of these structures, these two building blocks are difficult to assemble into a direct rod-connected diamond lattice, which is considered to be a champion photonic crystal. Here, we present a DNA origami-route for a direct rod-connected diamond photonic crystal exhibiting a complete photonic bandgap (PBG) in the visible regime. Using a combination of electromagnetic, phononic, and mechanical numerical analyses, we identify (i) the structural constraints of the 50 megadalton-scale giant DNA origami building blocks that could self-assemble into a direct rod-connected diamond lattice with high accuracy, and (ii) the elastic moduli that are essentials for maintaining lattice integrity in a buffer solution. A solution molding process could enable the transformation of the as-assembled DNA origami lattice into a porous silicon- or germanium-coated composite crystal with enhanced refractive index contrast, in that a champion relative bandwidth for the photonic bandgap (i.e., 0.29) could become possible even for a relatively low volume fraction (i.e., 16 vol%).

---

Since the pioneering works of E. Yablonovitch^1^ and S. John in 1987^2^, photonic crystals have transformed various fields including lasers, optomechanics, optoelectronics, and structural colorizations.^3–15^ According to on-demand applications, different manufacturing methods can be suitably chosen for their materializations. For example, serial-type monolithic lithography such as electron-beam or focused ion-beam lithography can be readily implemented for the fabrication of ultra-small lasers and optomechanical cavities because these applications depend on discrete structural units at the nanoscale.^3–8^ By contrast, structural colorization requiring large-area pixelation of visible photonic crystals is difficult to achieve with such serial-type fabrications.^9–15^

Instead, soft self-assembly has provided a viable route toward addressing scalability issues, due to this method being highly amenable for cost-effective solution processes.^9–14^ However, thus far, the building blocks for self-assembled visible photonic crystals have been mainly limited to colloidal spheres and block copolymers (BCPs), which in turn restricts the accessible 3D lattices to mainly face-centered cubic (FCC) for colloidal spheres and double gyroids (DG) for BCP. Unfortunately, neither FCC nor DG lattices are able to exhibit a complete PBG, which is essential for monotonic structural color independent of the viewing angle.^10–14^

Approximately 30 years ago, the diamond lattice was discovered as a champion photonic crystal due to enabling a complete and wide PBG with relatively low refractive index contrast between the host medium and lattice (*n*_cont_, ~ 0.8) and filling fraction (*f*lower than 0.5).^16,17^ In particular, a direct rod-connected diamond lattice can exhibit a wide and complete PBG, e.g., a relative bandwidth for photonic bandgap (*δ* = (Δω/ω)_max_) of ~ 0.3, with *f* of 0.19 and *n*_cont_ of 2.6.^17^ A modest lowering of *f* in photonic crystals is desired in terms of widening the PBG, since intuitively increasing the effective *n*_cont_; however, paradoxically, lowering *f* makes the fabrication of the 3D photonic crystals more challenging due to weakening the mechanical properties.^18^ This paradox becomes more pronounced for visible photonic crystals, where the scales of the primitive cell needs to be a few hundred nanometers. Indeed, previous reports exist only for experimental realizations of direct rod-connected diamond lattices that operate from the IR to microwave regimes.^19,20^ Even though the pyrochlore lattice was recently probed as a versatile-to-craft diamond visible photonic crystal by molecularly programmed self-assembly of colloidal spheres,^21–24^ the current theoretically exploited maximum *δ* is relatively small (e.g., ~ 0.20).^22–24^ These limitations on self-assembly originate from the restricted libraries of building blocks in conjunction with the relevant assembly techniques.

DNA can be molded into nearly any 2D and 3D shapes via molecular folding (i.e., DNA origami).^25–37^ Furthermore, it was found that DNA origami can be crystallized into 3D superlattices over large areas.^36,37^ As such, 3D crystallization of DNA origami could form superlattices, which have not been constructed thus far. This recent success in DNA nanotechnology leads us to raise the question: can we obtain a complete visible PBG with a lower *n*_cont_ and *f* by using a DNA-based, direct rod-connected diamond lattice? The designs for the DNA origami lattice and postmolding processes outlined in this work posit that *δ* of 0.17 ~ 0.29 can be achieved even with a relatively low *f* of 0.16 ~ 0.21 and *n*_cont_ of 2.0 ~ 3.0, demonstrating the technological viability of the DNA origami-route for diamond photonic crystals exhibiting a complete PBG in the visible regime.

Our aim in this work is to theoretically exploit the possibility of assembling of DNA origami into a direct rod-connected diamond photonic crystal that can exhibit a complete PBG in the visible regime. Toward this end, our work is organized as follows. First, we compare the accessible ranges of the PBG from two different diamond lattices that could be constructed by DNA origami self-assembly: (i) direct rod-connected (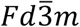, No. 227) and (ii) tetrahedral cage-connected lattices (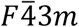, No. 216). Then, we design the 50 megadalton (MDa)-scale DNA origami building blocks that can be assembled into the target diamond lattice with a complete PBG in the visible regime. In particular, these DNA origami building blocks were designed to have sufficiently high mechanical stability at the junction point, which is foundational for high accuracy-assembly. Additionally, using the concept of elastic moduli, we quantify the mechanical stability of the assembled diamond lattice constructed by DNA origami because the structural integrity of the diamond lattice needs to be maintained during the postmolding process (i.e., decorating the DNA origami frame with a high refractive index material). Finally, we evolve the PBG of the DNA origami diamond lattice with respect to the postmolding process and confirm that the designed diamond lattice indeed exhibits a complete PBG in the visible regime.

In 2016, O. Gang et al. first reported that DNA origami can be crystallized into a diamond lattice.^36^ More specifically, the base-complementary binding between the tetrahedral DNA origami cage enables the crystallization of a diamond lattice of space group 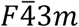, No. 216 (hereafter referred to as a tetrahedral cage-connected diamond lattice), as shown in **Figure 1a**. Notably, this DNA diamond lattice (the *n* of the DNA can be set at 1.8^38^) can open a complete PBG in air even with a relatively low *n*_cont_ of ~ 0.8 (see **Figure S1**, Supporting Information). In general, the diamond lattice requires a much lower *n*_cont_ to open a complete PBG compared to other lattices.^16,17^ However, as previously reported, a 3D lattice fully constructed by DNA origami is prone to structural deformations after the drying of the buffer solution.^39^ Molding the lattice with a 20 vol% higher refractive index and a stronger semiconductor (e.g., gallium arsenide (GaAs) with *n* of 3.6) can yield a mechanically stable 3D diamond lattice in air (*n*_cont_ of ~ 2.6) and open a wider complete PBG, as presented in **Figure 1b**. Nevertheless, the relative bandwidth of the complete PBG (*δ* ~ 0.20) is not comparable to that of the direct rod-connected counterpart containing the same vol% of GaAs (*δ* ~ 0.30) (see **Figures 1c-d**). Herein, a direct rod-connected diamond lattice can be assembled from tetrapod-shaped building blocks. Furthermore, the tetrahedral cage-connected diamond lattices previously reported had too small of a lattice constant (*a* of ~ 51 nm)^36^ and were unable to open a PBG in the visible regime. Therefore, we turned our attention to identifying what (other) DNA origami motifs need to be designed to obtain direct rod-connected diamond crystals with *a* of a few hundred nanometers and how to obtain a complete visible PBG by a postmolding process.

**Figure 1.**
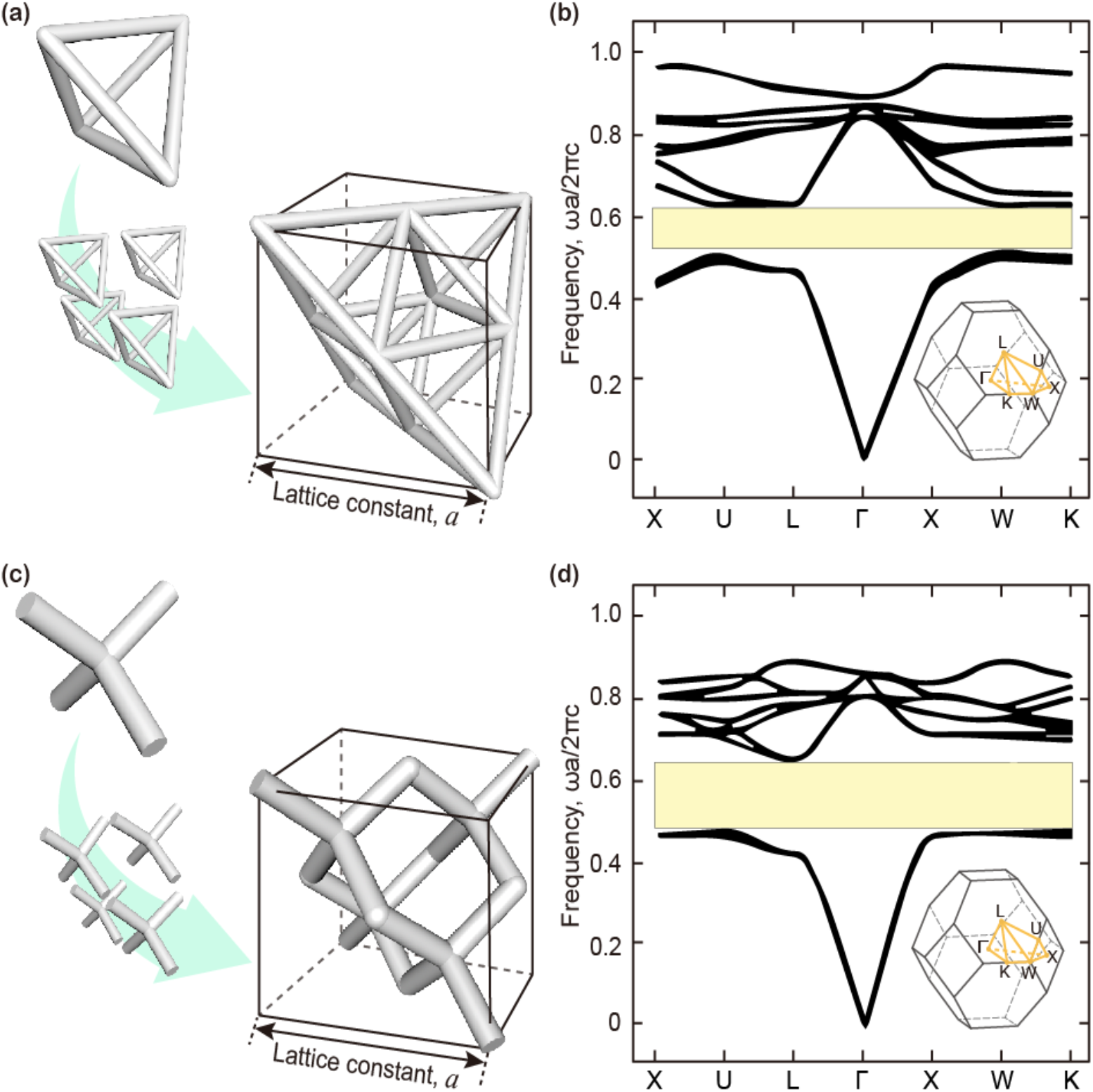
Diamond lattice, accessible with 3D DNA origami building blocks and photonic bandgap (PBG). (a) Tetrahedral cage-connected diamond lattice (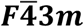, No. 216). (b) Corresponding PBG after molding the lattice with high refractive index (*n*) materials (e.g., gallium arsenide (GaAs) with *n* of 3.6). (c) Direct rod-connected diamond lattice (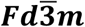, No. 227). (d) Corresponding PBG after molding the lattice with high *n* materials (e.g., GaAs with *n* of 3.6).

In 2018, T. Liedl et al. reported that DNA origami composed of the interconnected three 14-HB (helix bundles, indicating the number of DNA duplex in DNA origami bundles), hereafter referred to as three-axis tensegrity, can crystallize into a rod-type rhombohedral lattice by blunt-end stacking.^37^ Even though this rod-type rhombohedral lattice is unable to open a complete PBG (see **Figure S2**, Supporting Information), such success in the large-area crystallization of 3D DNA origami inspired us to design a direct rod-connected diamond lattice that could be assembled with base-complementary binding or blunt-end stacking between a rationally designed tetrapod of 3D DNA origami (**Figure 2**). Herein, a tetrapod of DNA origami consists of four equivalent arms with symmetric geometry; as shown in **Figure 2a**, the width and length of each arm are denoted *d* and *l*, respectively. To push the PBG wavelength to the visible regime, the corresponding *a* of the direct rod-connected diamond lattice should be at least 330 nm even under *n*_cont_ of 2, as systematically exploited in **Figure S3**, Supporting Information. Since the *l* of the tetrapod is geometrically set by 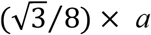, the four arms of the tetrapod must be at least ~ 70 nm in *l* for DNA origami building blocks.

**Figure 2.**
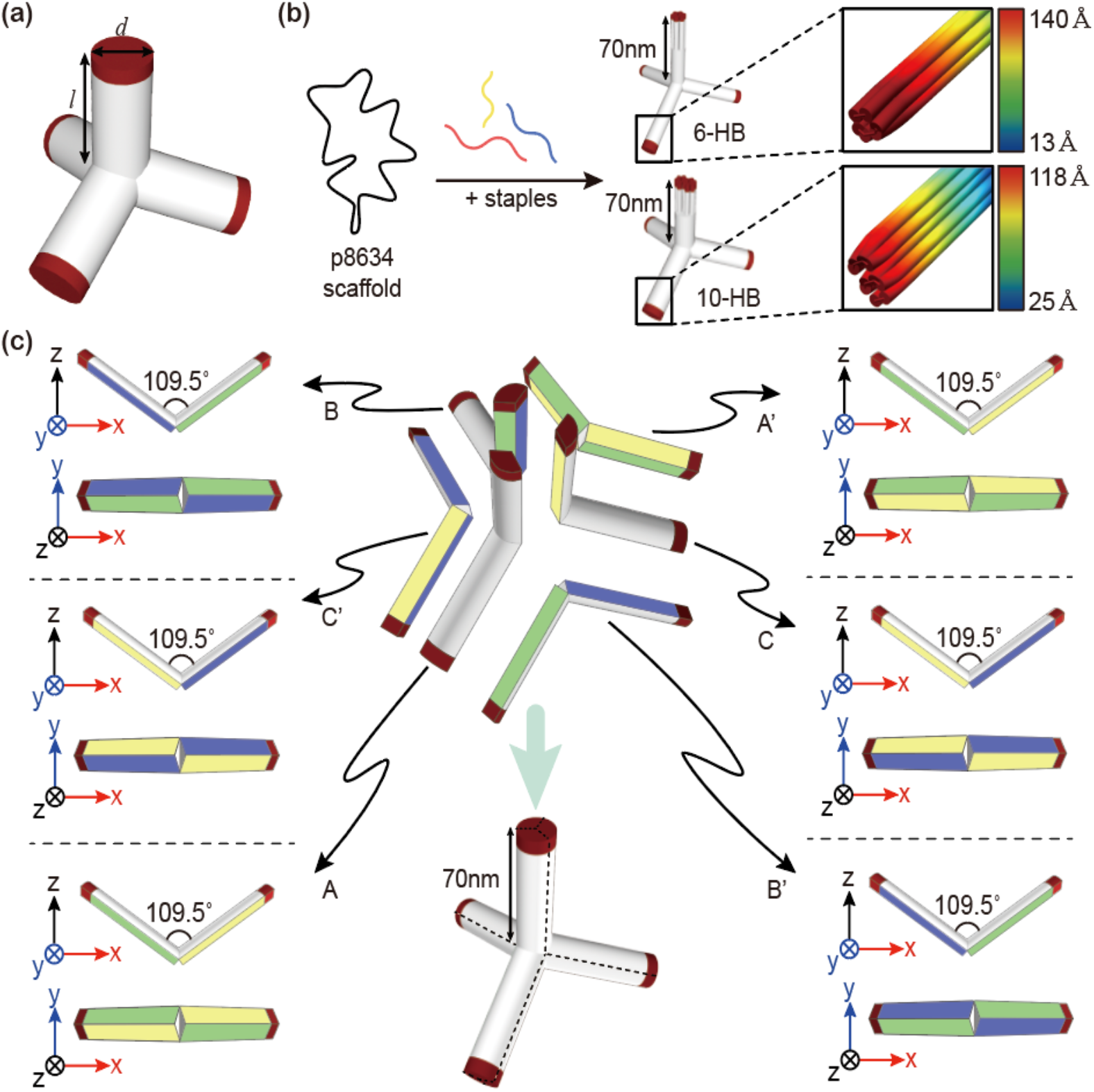
(a) Schematic for tetrapod that can be self-assembled into a direct rod-connected diamond lattice. The length and width of each arm of tetrapod are denoted *l* and *d*, respectively. (b) The maximum dimensions of the tetrapod that can be assembled by using a single scaffold (p8634) are 70 nm in *l* and 8~9 nm in *d* (10-HB). Given a honeycomb lattice, which is widely used for 3D DNA origami, 6- or 10-HB can be implemented into each arm of the tetrapod, self-assembled from a single scaffold. In these cases, the root mean square fluctuations (RMSF) of arm end, supported by CanDo simulation^42,43^ can be increased up to 140 Å. (c) The hierarchical assembly route for a giant DNA origami tetrapod (~ 50 MDa). Geometrically, the tetrapod can be divided into six equivalents with the shape of 109.5°-angled wings, highlighted by A/A’/B/B’/C/C’. Herein, the same false colored facets of each wing indicate the positions where sequence- or blunt-end stacking-based shape-complementary binding can occur.

Given the distance between the neighboring bases (i.e., 0.33 ~ 0.34 nm),^40^ the full folding of the longest scaffold DNA (i.e., p8634) allows us to obtain duplex DNA with 2849 ~ 2935 nm in total length (~ 8 MDa). Thus, each arm of the tetrapod can have up to 10-HB by full folding of single scaffold DNA, as shown in **Figure 2b** (i.e., 2935 nm/4 arms/70 nm = ~ 10.48). Considering the honeycomb lattice widely used in 3D DNA origami, the cross-sectional *d* can be ranged, for example, from 6 ~ 7 nm (for 6-HB) to 8 ~ 9 nm (for 10-HB). Thus, the corresponding aspect ratio (AR) of each tetrapod arm is greater than 7, which is much higher than those of DNA origami tetrahedron cages (cross-section of 10-HB and frame length of 40 nm; corresponding AR of ~ 4) and three-axis tensegrity (cross-section of 14-HB and frame length of 67 nm; corresponding AR of ~ 5), both of which were previously found to assemble well into diamond and rhombohedral lattices, respectively.^36,37^ Recently, it was found that the rigidity of DNA origami building blocks is the key to success in hierarchical assembly into superstructures or 3D crystals.^41^ Although the structural rigidity can vary not only due to the AR but also due to the overall geometry, a relatively high AR can make DNA origami too floppy in solution, implying that the assembly could be error-prone. The numerically simulated maximum root mean square fluctuations (RMSF) of the DNA origami tetrapods with cross-sections of 6- and 10-HB and *l* of 70 nm, which were supported by CanDo^42,43^, were found to increase to 118 Å ~ 140 Å at the ends of the arms, where base-complementary binding or blunt-end stacking should occur. The corresponding caDNAno designs^44^ that were used for the CanDo simulations^42,43^ are included in **Figure S4**, Supporting Information. These relatively high RMSF at the binding points should reduce the assembly accuracy, which in turn restricts the large-area 3D crystallization.^37,41^

To impart more mechanical rigidity to the DNA origami tetrapods, the number of HB in each arm should be further increased (i.e., AR should be reduced). Under the length constrains of each arm of the tetrapod building blocks (i.e., 70 nm), the AR cannot be reduced from 7 by the full folding of a single scaffold DNA, as mentioned above. To overcome the problem of the limited scaling up from a single scaffold, we conceived the hierarchical assembly of the DNA origami unit into a thicker-armed tetrapod, as presented in **Figure 2c**. The tetrapod can be divided into six equivalent subsets of 109.5°-angled wings with wing lengths of 70 nm, which are denoted A/A’/B/B’/C/C’ in **Figure 2c**. Three 109.5°-angled wings were designed to be assembled into half of a tetrapod (i.e., assemblies of A-B-C and A’-B’-C’); then, two halves of a tetrapod can be paired with each other, for example, by blunt-end shape-complementary binding.^31^ Thereby, each arm of the tetrapod was designed to consist of three wings.

The power of this hierarchical strategy lies in the versatile scale-up of the assembled tetrapod by using a single DNA origami motif. For example, each wing of the DNA origami with a cross-section of up to 20-HB and *l* of 70 nm can be obtained by a full folding of a single p8623 (8623*0.34 nm per base pair/2 arms of wings/70 nm spanning wing length = ~ 20.94 HB). Thereby, the number of HB in each arm of the assembled tetrapod could be widely tuned from 18 (6-HB per wing × 3) to 60 (20-HB per wing × 3).

In addition, we can use the folding of two scaffolds to further increase the number of HB in each arm of the tetrapod. As with the previous reports,^41^ we can use coassembly of p8064 and p7560 into a single DNA origami motif. In this case, 70 nm-long wings with much thicker cross-section (up to 37-HB) can be obtained. As a representative example, we designed 32-HB wings to be hierarchically assembled into tetrapods with cross-sections of 84-HB (*d* of ~ 25 nm and AR of 2.8) and *l* of 70 nm (**Figures 3a-c**). Herein, a total of 12-HB was designed to overlap between each wing, consequently providing the shape-complementary binding positions. The corresponding caDNAno designs are included in **Figures S5-S7**, Supporting Information. These thicker wings showed much smaller RMSF (up to 11 Å, as shown in **Figure 3a**), compared with those of 6- and 10-HB arm of tetrapods. The tetrapods self-assembled from six 32-HB wings (**Figure 3b**) can have cross-sections of 84-HB (**Figure 3c**), and the resultant RMSF at the binding points (ends of the tetrapod) was further reduced to 7 ~ 10 Å (**Figure 3d**). As such, the 3D crystallization of these giant tetrapods (50 MDa) into a direct rod-connected diamond lattice could be achieved with high accuracy (**Figure 3e**). We designed the ends of the assembled tetrapods to be nonflat for shape-complementary binding (i.e., programmable blunt-end stacking) to prevent defect came from mismatched blunt-end sites.^28,41^

**Figure 3.**
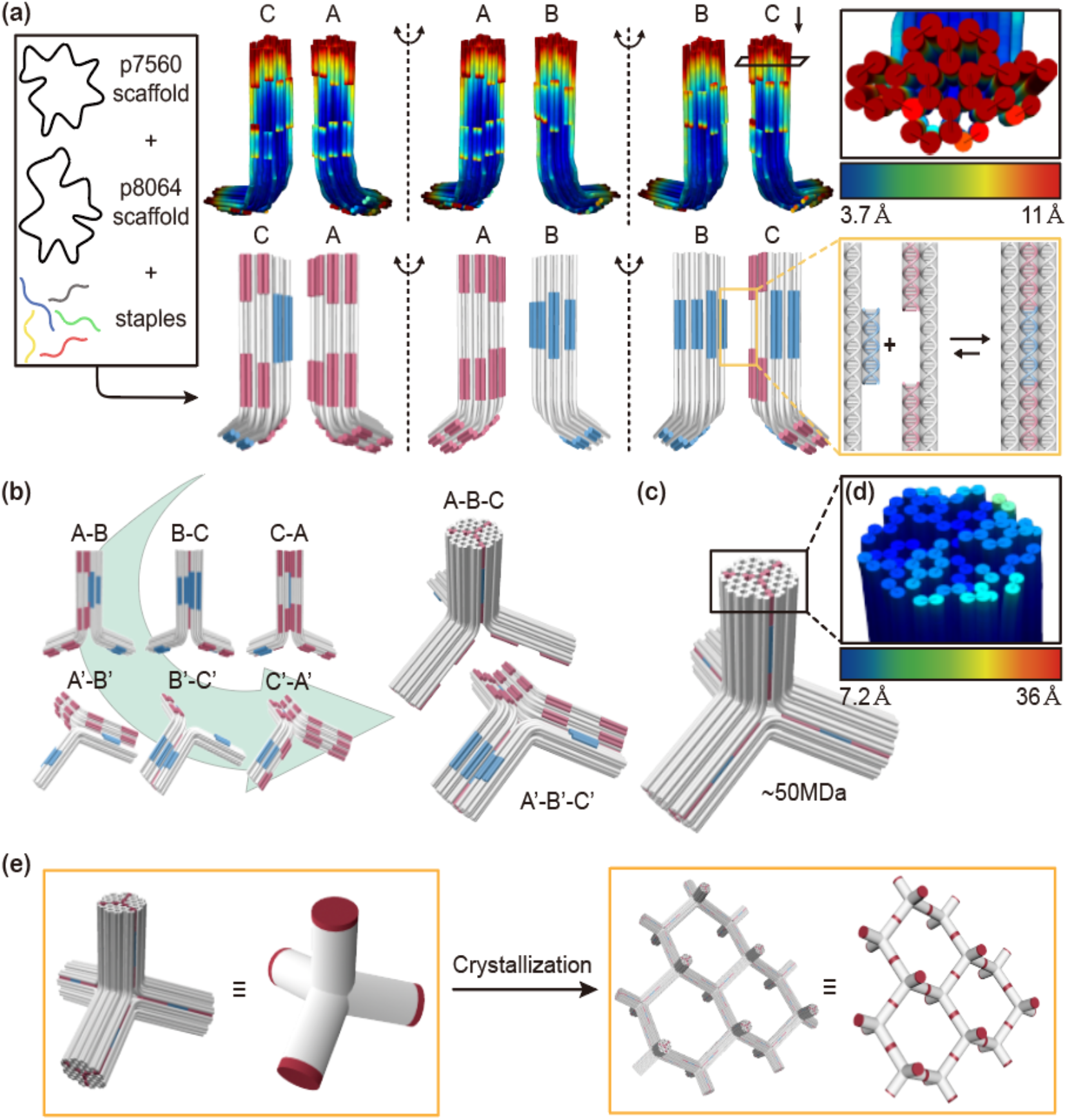
(a) 109.5°-angled wings with 32-HB that can be attained by coassembly of two scaffold strands (e.g., p7560 and p8064), corresponding RMSF obtained by CanDo simulation^42,43^, and route for shape-complementary binding between 109.5°-angled wings. RMSF of 109.5°-angled wing end was approximately 11 Å. (b) Assembling of half of the tetrapod (i.e., assemblies of A-B-C and A’-B’-C’). (c) Pairing of two halves of tetrapod by blunt-end shape-complementary binding. Then, a giant DNA origami tetrapod with 70 nm in *l* and 25 nm in *d* (84-HB) can be obtained. As a result, each arm of this tetrapod was designed to consist of three wings. (d) RMSF of each arm end of a giant DNA origami tetrapod, obtained by CanDo simulation^42,43^, was reduced to 7.2 Å. (e) Due to the low RMSF, a giant DNA origami tetrapod can crystallize into a direct rod-connected diamond lattice with a lattice constant (*a*) of 330 nm.

From an experimental perspective, the generalization of such DNA origami hierarchical assembly is still being elucidated. Thus far, the experimental verification of this method has been reported under very limited structural motifs such as small sized tetrahedron cages, three-axis tensegrities, and V-shaped multilayer objects.^36,37,41^ Advances in the thermodynamic analysis of DNA origami hierarchical assembly, possibly supported by molecular dynamics (MD) simulations, need to be followed; however, in contrast to other colloidal sciences,^45–47^ the efforts toward this theoretical understanding have not yet been validated in the field of DNA origami. In line with this, thermodynamic analysis, which could guide the optimal interaction conditions for high-yield hierarchical assembly of DNA origami, will be reported separately, being beyond the scope of this study.

The DNA origami diamond lattice, assembled within a buffer solution, needs to maintain structural integrity during the solution-based postmolding process. As already noted in **Figure S1**, the DNA origami diamond lattice in air (*n*_cont_ of 0.8) can open a complete PBG, but maintaining the lattice integrity of the 3D DNA origami crystals in air is not possible.^39^ Even though 3D DNA origami crystals can be highly stabile in a buffer solution, a complete PBG cannot be opened in this case due to the relatively low *n*_cont_ (~ 0.45). This problem can be addressed by a conformal coating of silica onto the DNA origami lattice in a buffer solution^39,48^ and subsequent molding of the lattice with high refractive index materials, as is detailed in the next section.

The lattice rigidity of the DNA origami diamond crystals in a buffer solution was quantified with the concept of effective elastic moduli, such as shear (*G*_eff_) and P-wave (*M*_eff_) moduli.^49,50^ Toward this end, we numerically simulated the phononic band structures of direct rod-connected DNA origami diamond lattices with same *a* (i.e., 330 nm) but different HB (i.e., 6, 24, and 84), as shown in **Figures 4a-c** (see details in Supporting Information). Then, we compared these phononic band structures with that of a three-axis tensegrity lattice (**Figure 4d**).^37^ In a previous report, the 3D crystal assembled from three-axis tensegrity DNA origami was found to maintain lattice integrity in a buffer solution well,^37^ serving as a good indicator for whether our DNA origami diamond lattice in a buffer solution could be maintained well or not.

**Figure 4.**
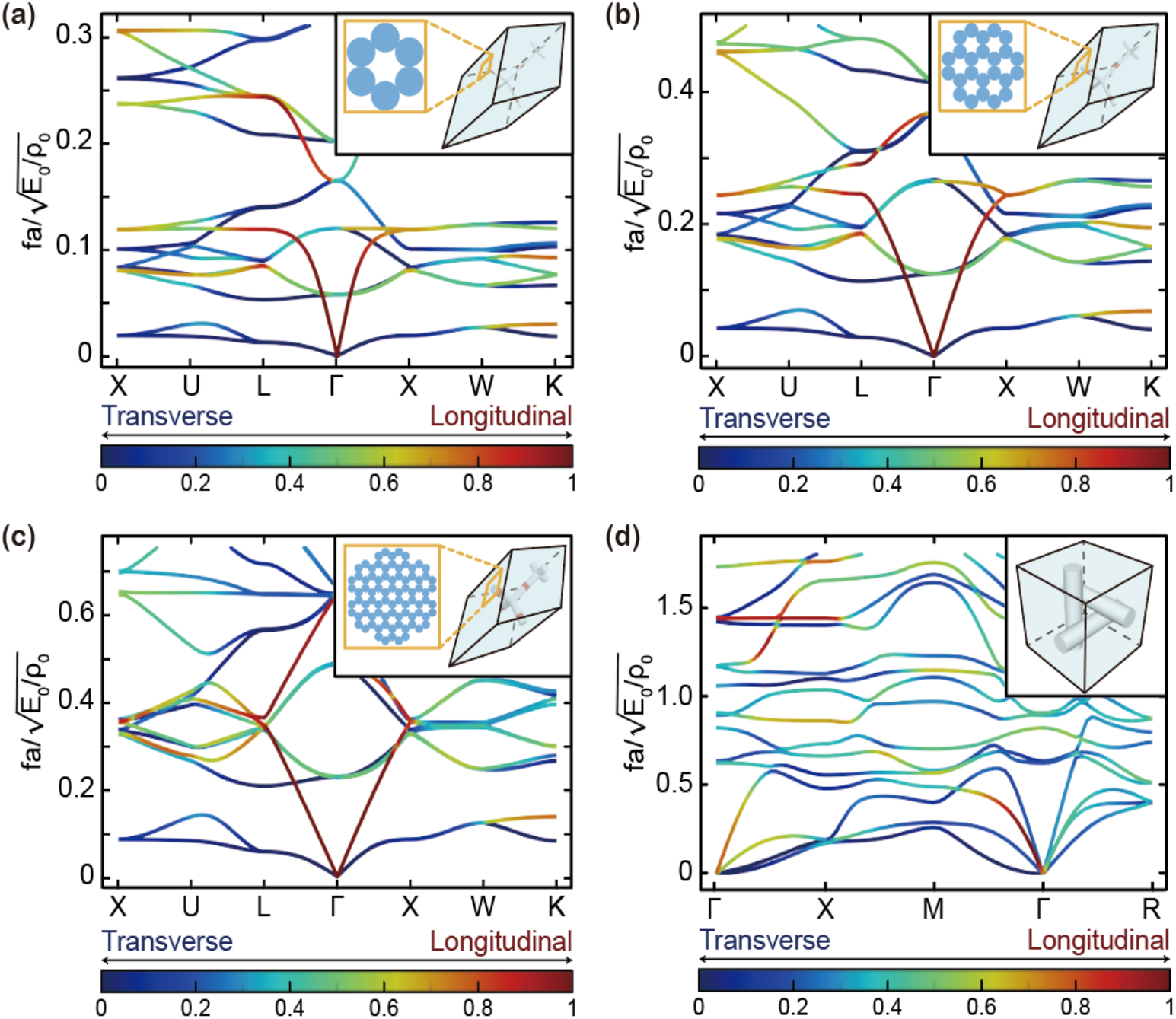
(a-d) Phonon band structures of the direct-rod connected (a) 6-HB, (b) 24-HB, (c) 84-HB diamond lattices and (d) three-axis tensegrity lattice.^37^ All the eigenfrequencies are non-dimensionalized with respect to the lattice constant (*a* = 330 nm), Young’s modulus (*E*_0_ = 300 MPa), and density (*ρ*_0_ = 1.6450 g/cm^3^). In (a), (b), and (c), the coordinates of the symmetry points in momentum space are as follows: X = [0, 0.5, 0.5], U = [1/4, 5/8, 5/8], L = [1/2, 1/2, 1/2], Γ = [0, 0, 0], W = [1/4, 3/4, 1/2], K = [3/8, 3/4, 3/8]. In (d), the coordinates of the symmetry points in momentum space are as follows: Γ = [0, 0, 0], X = [1/2, 0, 0], M = [1/2, 1/2, 0], R = [1/2, 1/2, 1/2].

In the phononic band structure of a 3D periodic array, generally, three bands spread from the Γ point. The two bands at the lowest frequencies and one band at a frequency higher than the lowest correspond to transverse and longitudinal waves, respectively (see **Figure 4**). The effective two transverse and one longitudinal wave velocities along the Γ-L and Γ-X (denoted *v*_*t*,1_, *v*_*t*,2_, and *v*_*l*_) were calculated by *ν* = *ω* / *k* around the Γ point. Once the input wavevector **k** was mapped in all directions in 3D space, the corresponding effective wave velocities were obtained in the same fashion. From the two effective transverse wave velocities obtained, the effective shear moduli were calculated by the relationships 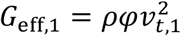 and 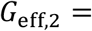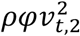, where *φ* is the volume fraction of the DNA origami. Similarly, the effective P-wave modulus was calculated from the relationship 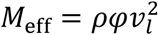. Finally, we profiled *G*_eff_ and *M*_eff_ with respect to the input wavevector **k** in all directions.

The collective sets of *G*_eff_ and *M*_eff_, obtained from our DNA origami diamond lattices and three-axis tensegrity lattice, are summarized in **Figures 5a-c**. The minimum *G*_1, eff_ (Min (*G*_1,eff_) of the direct rod-connected diamond lattices with 6- and 24-HB was found to be higher than that of the three-axis tensegrity lattice (**Figure 5a**). By contrast, Min (*G*_2,eff_) and Min (*M*_eff_) of the 6- and 24-HB diamond lattices were smaller than those of the three-axis tensegrity lattice (**Figures 5b-c**). Therefore, the 6- and 24-HB diamond lattices should be mechanically weaker than the three-axis tensegrity lattice, particularly along the directions of *G*_2,eff_ and *M*_eff_.

**Figure 5.**
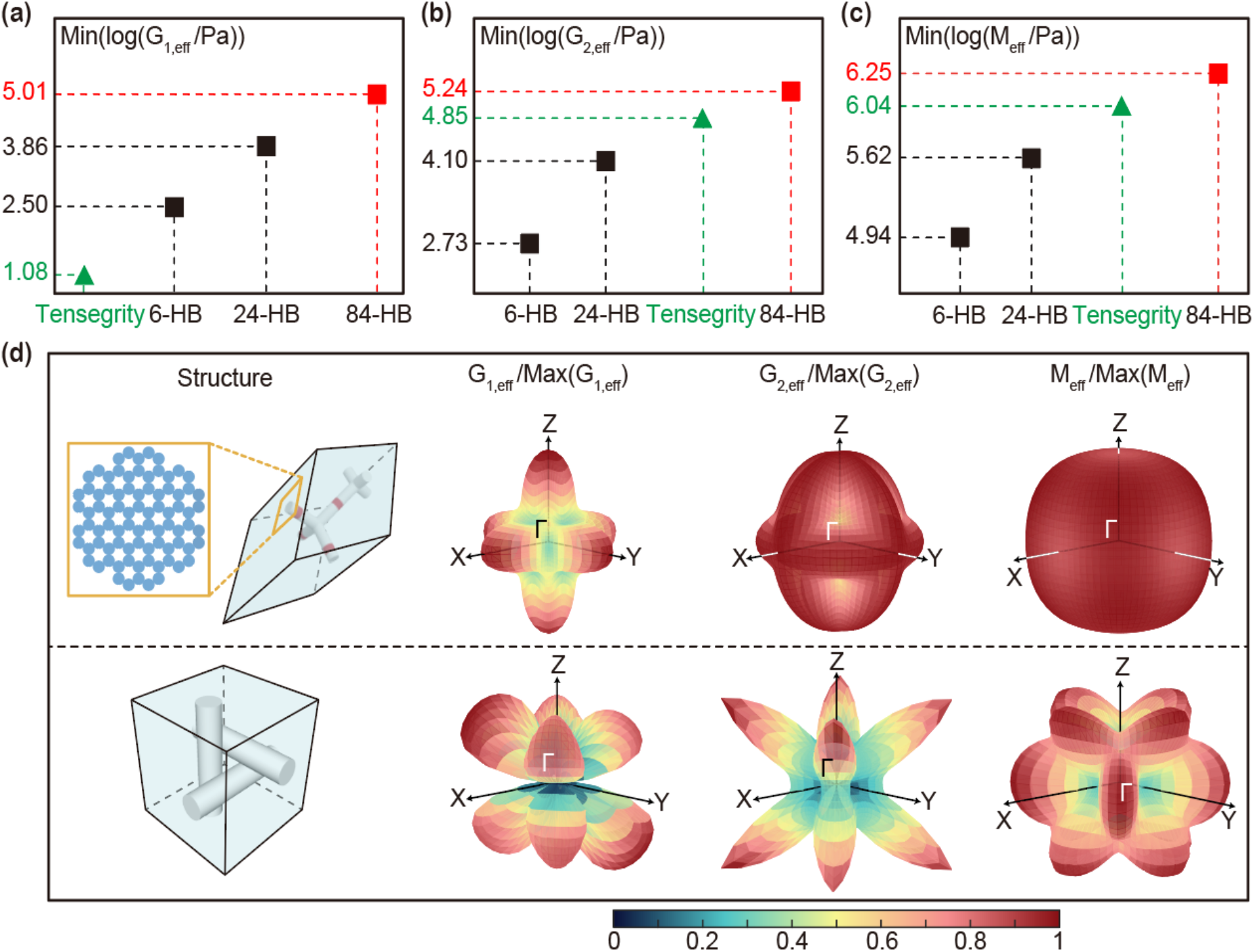
(a, b, c) Comparisons of shear (*G*_1,eff_ and *G*_2,eff_) and P-wave (*M*_eff_) moduli of DNA origami diamond lattices (Min (***G*_1,eff_**), Min (***G*_2,eff_**), and Min (***M*_eff_**)). Herein, the primitive cells together with the cross-sections of the three direct-rod diamond structures are depicted; the lattice vectors were a_1_=[0,1,1]*a*/2, a_2_=[1,0,1]*a*/2, and a_3_=[1,1,0]*a*/2. (d) Spatial distributions of ***G*_1,eff_**, and ***G*_2,eff_** and ***M*_eff_** for 84-HB direct rod-connected diamond lattice and three-axis tensegrity lattice.

One the one hand, the direct rod-connected diamond lattice with 84-HB exhibited higher Min (*G*_1,eff_), Min (*G*_2,eff_), and Min (*M*_eff_) than the three-axis tensegrity lattice (**Figures 5a-c**). Furthermore, the spatial distributions of *G*_1,eff_, *G*_2,eff_, and *M*_eff_ of this diamond lattice were more isotropic than those of the three-axis tensegrity lattice, as clearly shown in **Figure 5d**. In a 3D lattice with anisotropic spatial distribution of effective elastic moduli, stress can be locally concentrated in the direction of a relatively low effective elastic moduli. Overall, we can conclude that the direct rod-connected diamond lattice with 84-HB exhibits sufficient stiffness and mechanical properties to maintain high lattice fidelity in a buffer solution.

Finally, we profiled a PBG of the direct rod-connected diamond lattice, assembled from 84-HB tetrapods. **Figure 6** shows the corresponding photonic band diagram with respect to the postmolding procedures. In a buffer solution, the PBG of the as-assembled DNA origami lattice (*n* of 1.8; see system 1 in **Figure 6a**) can be partially opened only from Γ to L, as shown in **Figure 6b**, but this is not so visible owing to a low *n*_cont_ (0.47, as the *n* of a buffer can be simplified as 1.33). Although changing the host medium from a buffer solution to air by drying can widen a PBG via an increase in *n*_cont_, the 3D DNA origami crystals were found to be easily deformed in air, as mentioned above.^39^ Therefore, we needed to conceive a scheme for the postmolding process that can open a complete PBG by increasing *n*_cont_ while simultaneously imparting enough mechanical rigidity to the DNA lattice in both air and buffer solution.

**Figure 6.**
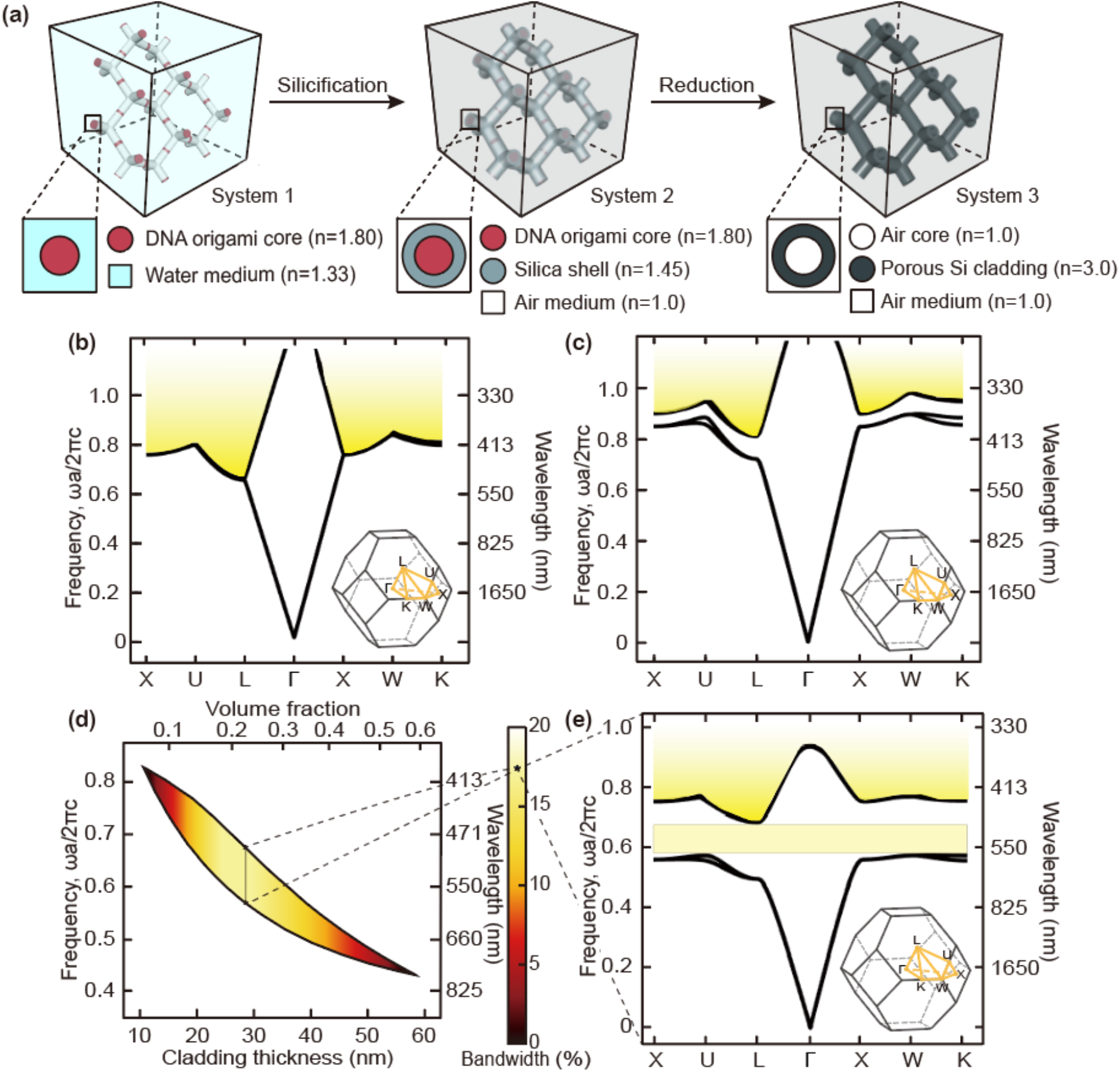
(a) Schematic for the postmolding process of a direct rod-connected diamond lattice. (b) Photonic band structure of as-assembled DNA origami lattice that is dispersed in a buffer solution (i.e., aqueous solution with salt ions). (c) Photonic band structure of DNA origami lattice that is conformably coated with 27 nm-thick silica (via silicification). Because the DNA origami lattice can be strengthened enough to maintain lattice integrity after silicification, a host medium of air can be justified. (d, e) Photonic band structure of the DNA origami lattice, coated with porous Si (Si with *n* of 3.0 and resultant *n*_cont_ of 2.0). The porous Si could be obtained by low-temperature reduction of silica with assistance from chemicals (e.g., magnesium).54,55 The relative bandwidth (*δ*) of the porous Si cladded DNA origami lattice varied according to Si thickness. A star (**\***) represents the maximum *δ* (i.e., 0.17 at a cladding thickness of 27 nm corresponding to *f* = ~ 0.21). A complete photonic bandgap is highlighted by a light yellow box. All the photonic band structures in Figure 6 were obtained by the finite element method (FEM) (see details in Supporting Information).^57^

Atomic layer deposition (ALD) or chemical vapor deposition (CVD) process, generally used for converting polymeric photonic crystal templates into high *n* counterparts,^20^, ^51–53^ are accompanied by vacuum conditions, which can readily deform 3D DNA origami crystals. Thereby, a solution-based process, by which the structural integrity of soft 3D DNA origami crystals can be maintained, needs to be implemented into the postmolding with a rigid material. Thanks to recent advances in the solution-based conformal coating of silica onto the surface of DNA origami (i.e., silicification in **Figure 6a**),^39,48^ this requirement can be effectively fulfilled. The stepwise reduction reactions consisting of prehydrolysing and cluster-induced silicification were found to conformably coat silica along the topology of the DNA origami.^39,48^ Furthermore, the thickness of this silica, which in turn defines *f*, can be deterministically adjusted according to the reaction time.^39,48^ As such, the DNA origami frame is strengthened via silicification in the buffer solution.

**Figure 6c** presents the photonic band diagram of the direct rod-connected diamond lattice composed of a DNA origami core and silica shell (i.e., system 2). Herein, the silica shell is assumed to be 30 nm thick (i.e., 21 vol%); we exploited this configuration only in an air host medium, as the lattice integrity of silica-strengthened DNA origami crystal can be maintained even in air. The PBG was not opened completely because the *n* of silica (~ 1.45) and the resultant *n*_cont_ (~ 0.45) are insufficient.

Recently, it turned out that magnesium or aluminum reduction at high temperature (i.e., 700°C) can chemically transform low *n* silica to high *n* silicon (Si).^54,55^ This method converts silica into porous rather than solid Si due to the reduction-mediated vaporization of oxygen, so as to limit the achievable *n* to 3.0, which is lower than the *n* of pure Si (4.0).^56^ Additionally, the area initially corresponding to the DNA origami frame should be converted into an air void (*d* of ~ 25 nm) during such a high temperature treatment. Given these realistic conditions (i.e., air core/porous Si cladding with *n*_cont_ of 2.0), we investigated the photonic band diagram according to the *f* of porous Si (i.e., system 3). As shown in **Figures 6d-e**, a complete visible PBG at wavelengths of 482~574 nm can be optimally opened with a *δ* of 0.17. The required vol% for this complete visible PBG was relatively small (21 vol%). This achievable *δ* in the visible regime can be further widen to 0.19, when *a* is increased to 380 nm (see **Figure S3**, Supporting Information). A coassembly of p8064 and p7560 can increase *a* up to 380 nm, once cross-section of a giant tetrapod is fixed to 84-HB. A detailed pipeline for the optimization of PBG and *δ* is identified in **Figure S3**, Supporting Information.

The silica-coated 3D DNA origami (i.e., system 2), which could be stable in air, becomes compatible with vacuum-based processes for higher *n* material deposition (*i.e.*, pure Si or germanium with *n* of 4) including ALD and CVD (see **Figure 7a**). Therefore, *n*_cont_ can be further increased to 3.0. **Figures 7b-c** show the possible PBG of the 84-HB DNA origami-based, direct rod-connected diamond lattice (*a* of 330 nm), consisting of air core/pure Ge cladding (i.e., system 4). According to the thickness (or vol%) of Ge cladding, *δ* was found to optimally increase to 0.26 and could be further widen (up to ~ 0.29) according to the lattice constant (**Figure S3**, Supporting Information). As summarized in **Figure 8**, our DNA origami-route for direct rod-connected diamond lattices allows achievement of relatively high *δ* with relatively low *f* compared with other self-assembled diamond lattices that have been theoretically suggested thus far (i.e., direct and indirect pyrochlore and diamond lattices).^22–24^

**Figure 7.**
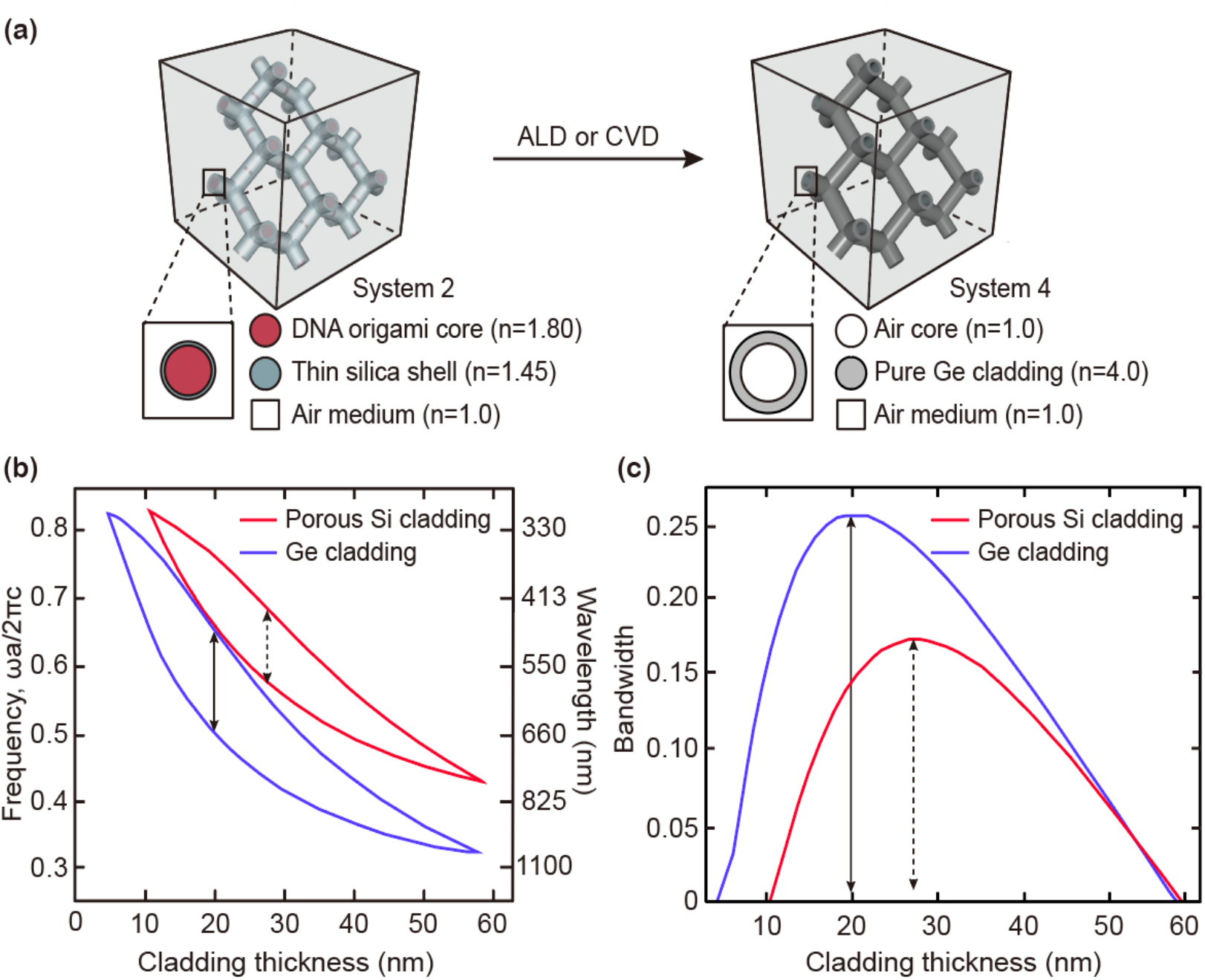
(a) Schematic of the CVD and ALD molding process of a direct rod-connected diamond lattice. Pure germanium (Ge) or Si with an *n* of 4.0 can be used as cladding. (b, c) Comparisons between photonic band structures of the DNA origami lattices coated with porous Si and pure Ge.

**Figure 8.**
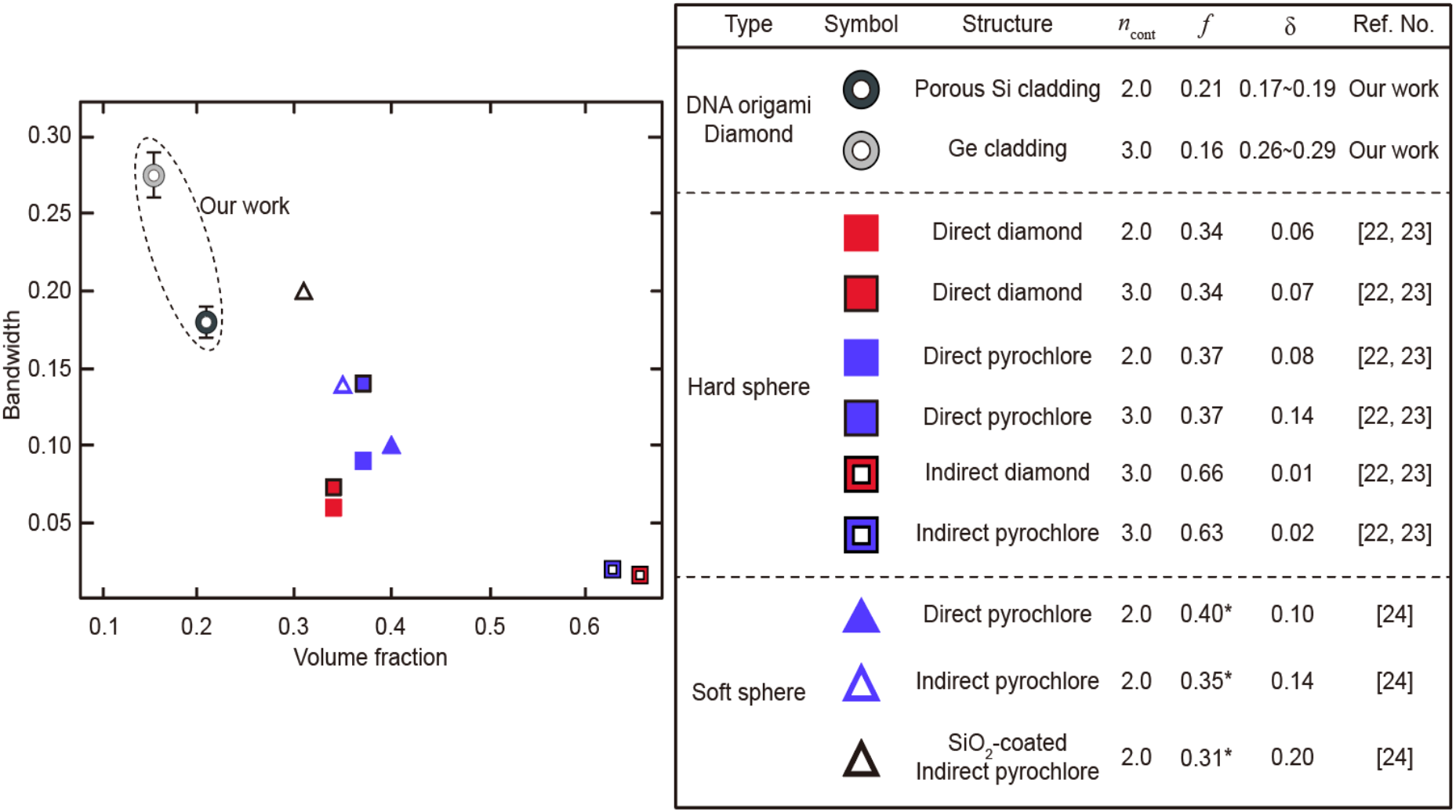
Comparison between maximum *δ* of our structures and those of other self-assembled diamond lattices suggested thus far (i.e., both (direct and indirect) pyrochlore and diamond lattices). Asterisks in Figure 8 (*) corresponding to the *f* are the values abstracted from structural information the literature.^22–24^

The power of 3D DNA origami photonic crystals lies in that this material can self-assemble into the nearly arbitrarily shaped, nanoscale building blocks. Using the design outlined in this work as a starting point, the 3D crystallization of DNA origami can greatly expand the accessible range of the photonic crystal lattices, which in turn enriches the engineering of PBGs in the visible regime. However, more in-depth follow-up verifications of numerical simulations (e.g., MD) are required to rigorously predict both the hierarchical assembly of DNA origami into a giant structure (i.e., 50 MDa tetrapods) and the 3D crystallization of this giant DNA origami. At the moment, development of appropriate computational methods in this direction remains elusive. Additionally, the aim of DNA origami photonic crystals is not limited to achieving a champion PBG. As the surface of these materials could be deterministically encoded with a variety of functionalities with molecular precision (e.g., quantum dots, fluorophores, and plasmonic nanoparticles), 3D DNA origami photonic crystals can be transformed into a general nanophotonic pegboard advancing the molding of light flow and solar energy harvesting.

## Supporting information

Supplemental information will be used for the link to the file on the preprint site

## Acknowledgement

This work was supported by Samsung Research Funding & Incubation Center for Future Technology of Samsung Electronics (Project Number SRFC-MA1801-04). S. H. Park acknowledges the scholarship support from the KU-KIST school project.

## Supporting Information

Details about (i) calculation of effective elastic moduli and photonic band structures, (ii) pipeline for optimal bandgap engineering, and (iii) design diagram for simulation units in manuscripts using software caDNAno ver 2.2.0^44^ are available in Supporting Information.

## References

(1) Yablonovitch, E., Inhibited Spontaneous Emission in Solid-State Physics and Electronics. Phys. Rev. Lett. 1987, 58, 2059–2062.

(2) John, S., Strong Localization of Photons in Certain Disordered Dielectric Superlattices. Phys. Rev. Lett. 1987, 58, 2486–2489.

(3) Campbell, M.; Sharp, D. N.; Harrison, M. T.; Denning, R. G.; Turberfield, A. J., Fabrication of Photonic Crystals for the Visible Spectrum by Holographic Lithography. Nature 2000, 404, 53–56.

(4) Park, H.-G.; Kim, S.-H.; Kwon, S.-H.; Ju, Y.-G.; Yang, J.-K.; Baek, J.-H.; Kim, S.-B.; Lee, Y.-H., Electrically Driven Single-Cell Photonic Crystal Laser. Science 2004, 305, 1444.

(5) Notomi, M.; Taniyama, H.; Mitsugi, S.; Kuramochi, E., Optomechanical Wavelength and Energy Conversion in High-Double-Layer Cavities of Photonic Crystal Slabs. Phys. Rev. Lett. 2006, 97, 023903.

(6) Eichenfield, M.; Camacho, R.; Chan, J.; Vahala, K. J.; Painter, O., A Picogram- and Nanometre-Scale Photonic-Crystal Optomechanical Cavity. Nature 2009, 459, 550–555.

(7) Eichenfield, M.; Chan, J.; Camacho, R. M.; Vahala, K. J.; Painter, O., Optomechanical Crystals. Nature 2009, 462, 78–82.

(8) Mak, K. F.; Shan, J., Photonics and Optoelectronics of 2D Semiconductor Transition Metal Dichalcogenides. Nat. Photonics 2016, 10, 216.

(9) Kim, H.; Ge, J.; Kim, J.; Choi, S.-E.; Lee, H.; Lee, H.; Park, W.; Yin, Y.; Kwon, S., Structural Colour Printing Using a Magnetically Tunable and Lithographically Fixable Photonic Crystal. Nat. Photonics 2009, 3, 534–540.

(10) Wang, J.; Wen, Y.; Ge, H.; Sun, Z.; Zheng, Y.; Song, Y.; Jiang, L., Simple Fabrication of Full Color Colloidal Crystal Films with Tough Mechanical Strength. Macromol. Chem. Phys. 2006, 207, 596–604.

(11) Urade, V. N.; Wei, T.-C.; Tate, M. P.; Kowalski, J. D.; Hillhouse, H. W., Nanofabrication of Double-Gyroid Thin Films. Chem. Mater. 2007, 19, 768–777.

(12) Arsenault, A. C.; Puzzo, D. P.; Manners, I.; Ozin, G. A., Photonic-Crystal Full-Colour Displays. Nat. Photonics 2007, 1, 468–472.

(13) Ahn, H.; Park, S.; Kim, S.-W.; Yoo, P. J.; Ryu, D. Y.; Russell, T. P., Nanoporous Block Copolymer Membranes for Ultrafiltration: A Simple Approach to Size Tunability. ACS Nano 2014, 8, 11745–11752.

(14) Lee, S. Y.; Kim, S. H.; Hwang, H.; Sim, J. Y.; Yang, S. M., Controlled Pixelation of Inverse Opaline Structures Towards Reflection-Mode Displays. Adv. Mater. 2014, 26, 2391–2397.

(15) Nam, S. H.; Park, J.; Jeon, S., Rapid and Large-Scale Fabrication of Full Color Woodpile Photonic Crystals via Interference from a Conformal Multilevel Phase Mask. Adv. Funct. Mater. 2019, 1904971.

(16) Ho, K. M.; Chan, C. T.; Soukoulis, C. M., Existence of a Photonic Gap in Periodic Dielectric Structures. Phys. Rev. Lett. 1990, 65, 3152–3155.

(17) Chan, C. T.; Ho, K. M.; Soukoulis, C. M., Photonic Band Gaps in Experimentally Realizable Periodic Dielectric Structures. Europhys. Lett. 1991, 16, 563–568.

(18) Maldovan, M.; Thomas, E. L., Diamond-Structured Photonic Crystals. Nat. Mater. 2004, 3, 593–600.

(19) Kirihara, S.; Takeda, M.; Sakoda, K.; Miyamoto, Y., Electromagnetic Wave Control of Ceramic/Resin Photonic Crystals with Diamond Structure. Sci. Technol. Adv. Mat. 2004, 5, 225–230.

(20) Chen, L., et al., Observation of Complete Photonic Bandgap in Low Refractive Index Contrast Inverse Rod-Connected Diamond Structured Chalcogenides. ACS Photonics 2019, 6, 1248–1254.

(21) Ducrot, É.; He, M.; Yi, G.-R.; Pine, D. J., Colloidal Alloys with Preassembled Clusters and Spheres. Nat. Mater. 2017, 16, 652.

(22) Hynninen, A.-P.; Thijssen, J. H. J.; Vermolen, E. C. M.; Dijkstra, M.; van Blaaderen, A., Self-Assembly Route for Photonic Crystals with a Bandgap in the Visible Region. Nat. Mater. 2007, 6, 202–205.

(23) Vermolen, E. C. M.; Thijssen, J. H. J.; Moroz, A.; Megens, M.; Blaaderen, A. v., Comparing Photonic Band Structure Calculation Methods for Diamond and Pyrochlore Crystals. Opt. Express 2009, 17, 6952–6961.

(24) Ducrot, É.; Gales, J.; Yi, G.-R.; Pine, D. J., Pyrochlore Lattice, Self-Assembly and Photonic Band Gap Optimizations. Opt. Express 2018, 26, 30052–30060.

(25) Rothemund, P. W. K., Folding DNA to Create Nanoscale Shapes and Patterns. Nature 2006, 440, 297–302.

(26) Ke, Y.; Sharma, J.; Liu, M.; Jahn, K.; Liu, Y.; Yan, H., Scaffolded DNA Origami of a DNA Tetrahedron Molecular Container. Nano Lett. 2009, 9, 2445–2447.

(27) Douglas, S. M.; Dietz, H.; Liedl, T.; Högberg, B.; Graf, F.; Shih, W. M., Self-Assembly of DNA into Nanoscale Three-Dimensional Shapes. Nature 2009, 459, 414.

(28) Woo, S.; Rothemund, P. W., Programmable Molecular Recognition Based on the Geometry of DNA Nanostructures. Nat. Chem. 2011, 3, 620–627.

(29) Han, D.; Pal, S.; Nangreave, J.; Deng, Z.; Liu, Y.; Yan, H., DNA Origami with Complex Curvatures in Three-Dimensional Space. Science 2011, 332, 342.

(30) Iinuma, R.; Ke, Y.; Jungmann, R.; Schlichthaerle, T.; Woehrstein, J. B.; Yin, P., Polyhedra Self-Assembled from DNA Tripods and Characterized with 3D DNA-PAINT. Science 2014, 344, 65.

(31) Gerling, T.; Wagenbauer, K. F.; Neuner, A. M.; Dietz, H., Dynamic DNA Devices and Assemblies Formed by Shape-Complementary, Non–Base Pairing 3D Components. Science 2015, 347, 1446–1452.

(32) Benson, E.; Mohammed, A.; Gardell, J.; Masich, S.; Czeizler, E.; Orponen, P.; Högberg, B., DNA Rendering of Polyhedral Meshes at the Nanoscale. Nature 2015, 523, 441–444.

(33) Veneziano, R.; Ratanalert, S.; Zhang, K.; Zhang, F.; Yan, H.; Chiu, W.; Bathe, M., Designer Nanoscale DNA Assemblies Programmed from the Top Down. Science 2016, 352, 1534.

(34) Linko, V.; Kostiainen, M. A., Automated Design of DNA Origami. Nat. Biotechnol. 2016, 34, 826–827.

(35) Nummelin, S.; Kommeri, J.; Kostiainen, M. A.; Linko, V., Evolution of Structural DNA Nanotechnology. Adv. Mater. 2018, 30, e1703721.

(36) Liu, W.; Tagawa, M.; Xin, H. L.; Wang, T.; Emamy, H.; Li, H.; Yager, K. G.; Starr, F. W.; Tkachenko, A. V.; Gang, O., Diamond Family of Nanoparticle Superlattices. Science 2016, 351, 582.

(37) Zhang, T.; Hartl, C.; Frank, K.; Heuer-Jungemann, A.; Fischer, S.; Nickels, P. C.; Nickel, B.; Liedl, T., 3D DNA Origami Crystals. Adv. Mater. 2018, 30, 1800273.

(38) Inagaki, T.; Hamm, R. N.; Arakawa, E. T.; Painter, L. R., Optical and Dielectric Properties of DNA in the Extreme Ultraviolet. J. Chem. Phys. 1974, 61, 4246–4250.

(39) Nguyen, L.; Doblinger, M.; Liedl, T.; Heuer-Jungemann, A., DNA-Origami-Templated Silica Growth by Sol-Gel Chemistry. Angew. Chem. Int. Ed. 2019, 58, 912–916.

(40) Saenger, W., Principles of Nucleic Acid Structure, Cantor, C. R., Eds. Springer 1984; pp 253–281.

(41) Wagenbauer, K. F.; Sigl, C.; Dietz, H., Gigadalton-Scale Shape-Programmable DNA Assemblies. Nature 2017, 552, 78.

(42) Castro, C. E.; Kilchherr, F.; Kim, D.-N.; Shiao, E. L.; Wauer, T.; Wortmann, P.; Bathe, M.; Dietz, H., A Primer to Scaffolded DNA Origami. Nat. Methods 2011, 8, 221–229.

(43) Kim, D.-N.; Kilchherr, F.; Dietz, H.; Bathe, M., Quantitative Prediction of 3D Solution Shape and Flexibility of Nucleic Acid Nanostructures. Nucleic Acids Res 2011, 40, 2862–2868.

(44) Douglas, S. M.; Marblestone, A. H.; Teerapittayanon, S.; Vazquez, A.; Church, G. M.; Shih, W. M., Rapid Prototyping of 3D DNA-Origami Shapes with caDNAno. Nucleic Acids Res 2009, 37, 5001–5006.

(45) Zhang, Z.; Glotzer, S. C., Self-Assembly of Patchy Particles. Nano Lett. 2004, 4, 1407–1413.

(46) Glotzer, S. C.; Solomon, M. J., Anisotropy of Building Blocks and Their Assembly into Complex Structures. Nat. Mater. 2007, 6, 557–562.

(47) Damasceno, P. F.; Engel, M.; Glotzer, S. C., Predictive Self-Assembly of Polyhedra into Complex Structures. Science 2012, 337, 453.

(48) Liu, X., et al., Complex Silica Composite Nanomaterials Templated with DNA Origami. Nature 2018, 559, 593–598.

(49) Mao, X.; Chen, Q.; Granick, S., Entropy Favours Open Colloidal Lattices. Nat. Mater. 2013, 12, 217.

(50) Hur, K.; Hennig, R. G.; Wiesner, U., Exploring Periodic Bicontinuous Cubic Network Structures with Complete Phononic Bandgaps. J. Phys. Chem. C 2017, 121, 22347–22352.

(51) King, J. S.; Graugnard, E.; Summers, C. J., Tio2 Inverse Opals Fabricated Using Low-Temperature Atomic Layer Deposition. Adv. Mater. 2005, 17, 1010–1013.

(52) Garcia-Santamaria, F.; Xu, M. J.; Lousse, V.; Fan, S. H.; Braun, P. V.; Lewis, J. A., A Germanium Inverse Woodpile Structure with a Large Photonic Band Gap. Adv. Mater. 2007, 19, 1567–1570.

(53) Hermatschweiler, M.; Ledermann, A.; Ozin, G. A.; Wegener, M.; von Freymann, G., Fabrication of Silicon Inverse Woodpile Photonic Crystals. Adv. Funct. Mater. 2007, 17, 2273–2277.

(54) Richman, E. K.; Kang, C. B.; Brezesinski, T.; Tolbert, S. H., Ordered Mesoporous Silicon through Magnesium Reduction of Polymer Templated Silica Thin Films. Nano Lett. 2008, 8, 3075–3079.

(55) Lai, Y.; Thompson, J. R.; Dasog, M., Metallothermic Reduction of Silica Nanoparticles to Porous Silicon for Drug Delivery Using New and Existing Reductants. Chem-Eur. J. 2018, 24, 7913–7920.

(56) Sohn, H., Handbook of Porous Silicon, Canham, L., Eds. Springer 2014; pp 341–352.

(57) Maldovan, M.; Thomas, E. L.; Periodic Materials and Interference Lithography, Wiley-VCH 2009; pp 141–181.

