## Supplemental information will be used for the link to the file on the preprint site for "Design of DNA Origami Diamond Photonic Crystals"

Table of Contents:

1. **Calculation of effective elastic moduli**
2. **Calculation of photonic band structures**
3. **Origami design and simulation**
4. **Photonic band structure for tetrahedron cage-connected diamond lattice full made of DNA**
5. **Photonic band structure for three-axis tensegrity**
6. **The pipeline for optimal bandgap engineering and corresponding results**
7. **Design diagram for simulation units**

### 1. Calculation of effective elastic moduli

To compute effective elastic moduli of the 3D DNA-origami arrays, at first, their phonon band structures are needed. For the given unit cell such as **Figure 4a-d**, following eigenfrequency equation with Floquet boundary conditions were computed:

$$-\rho\omega^2\mathbf{u} - \nabla\cdot\boldsymbol{\sigma} = \mathbf{F}_v \quad \text{(S1)}$$

$$\mathbf{u}_2 = \mathbf{u}_1 e^{i\mathbf{k}_f\cdot(\mathbf{r}_2-\mathbf{r}_1)} \quad \text{(S2)}$$

where **u** is the displacement vector as the eigenstates, $\rho$ is the density of the DNA origami, $\omega$ is the eigenfrequency (angular frequency, $\omega = 2\pi f$), $\sigma$ is the Cauchy stress, and $\mathbf{F}_v$ is the volume force. $\mathbf{k}_f$ and **r** mean the wave vector and position vector, respectively. Herein, subscript 1 and 2 indicate the indices of boundaries which define the periodicity of the 3D structure. As with the design in manuscript, the lattice constant was set by 330 nm. The materials properties of DNA duplex, used in this calculation, were as follows: Young's modulus of 300 MPa, Poisson's ratio of 0.48, and the density of 1.6450 g/cm$^3$. All the simulations were carried out using commercial finite element method (FEM).

To identify the nature of the phonon wave, the displacement vector was written as $\mathbf{u} = \mathbf{u}_L + \mathbf{u}_T$ where $\mathbf{u}_L$ and $\mathbf{u}_T$ are its longitudinal and transverse components, respectively. The longitudinal component was expressed as $\mathbf{u}_L = (\mathbf{k}\cdot\mathbf{u}/||\mathbf{k}||^2)\,\mathbf{k}$, while the transverse component was given by $\mathbf{u}_T = \mathbf{u} - (\mathbf{k}\cdot\mathbf{u}/||\mathbf{k}||^2)\mathbf{k}$. The ratio of longitudinal component to transverse component was then written as $\langle\mathbf{u}_L|\mathbf{u}_L\rangle/(\langle\mathbf{u}_L|\mathbf{u}_L\rangle + \langle\mathbf{u}_T|\mathbf{u}_T\rangle)$, where $\langle\mathbf{a}|\mathbf{b}\rangle$ corresponds to $\int_{cell}(\mathbf{a}^*\cdot\mathbf{b})d\mathbf{r}^3$. This ratio was quantitated as the reddish to bluish colors, as presented in **Figures 4a-d**.[1]

### 2. Calculation of photonic band structures

All photonic band structures shown in this study were computed using following eigenfrequency equation with Floquet boundary conditions:

$$\nabla\times\left(\frac{1}{\mu_r}\nabla\times\mathbf{E}\right) = \omega^2\frac{\varepsilon_r}{c^2}\mathbf{E} \quad \text{(S3)}$$

$$\mathbf{E}_2 = \mathbf{E}_1 e^{i\mathbf{k}_f\cdot(\mathbf{r}_2-\mathbf{r}_1)} \quad \text{(S4)}$$

where **E** is the electric field as the eigenstates, $c$ is the speed of light in vacuum, $\varepsilon_r(\mathbf{r})$ and $\mu_r(\mathbf{r})$ are the relative permittivity and permeability of the structure with respect to the position **r**, and $\omega$ is the eigenfrequency (angular frequency, $\omega = 2\pi f$). $\mathbf{k}_f$ and **r** indicate the wave vector and position vector, respectively, and subscripts 1 and 2 correspond to the indices of boundaries, which define the periodicity of the 3D structure. We assumed $\mu_r$=1 for all cases. Refractive index $n(\mathbf{r})$ was input through $\varepsilon_r(\mathbf{r})$, i.e., $\varepsilon_r(\mathbf{r}) = \{n(\mathbf{r})\}^2$; $n$ of the materials used in this study include **Re** and **Im** parts. All the simulations were carried out using FEM.

#### 3. Origami design and simulation

DNA origami designs for the direct-rod connected diamond lattices with 6-HB, 10-HB, and 32-HB were performed by using caDNAno version 2.2.0.[2] Due to its generality, honeycomb lattice was chosen as 3D lattice geometry. Then, the pre-designed caDNAno files were used for the numerical predictions of their mechanical properties (i.e., maximum root mean square fluctuations (RMSF)) by using CanDo.[3,4] We precisely tuned bp/turn and helix diameter to minimize structural distortions; 10.34 bp/turn with 2.5 nm of each helix diameter was optimized condition.

#### 4. Photonic band structure for tetrahedron cage-connected diamond lattice full made of DNA

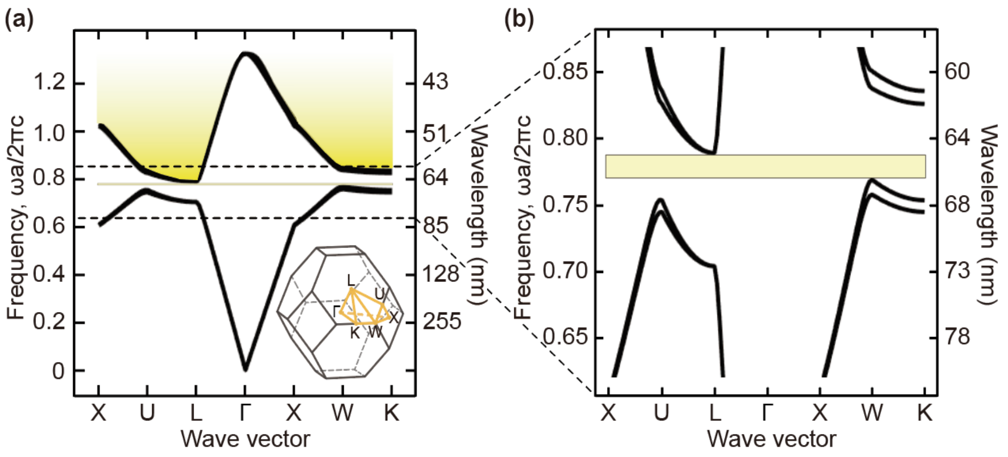


**Figure S1.** (a) Photonic band structure for the assembled tetrahedron cage with pure DNA origami in air medium ($n_{cont}$ ~ 0.8). (b) Magnified view for a complete PBG. All eigenfrequencies were non-dimensionalized with respect to the lattice constant ($a$ = 51 nm). The host medium is assumed to be air.

5. Photonic band structure for three-axis tensegrity

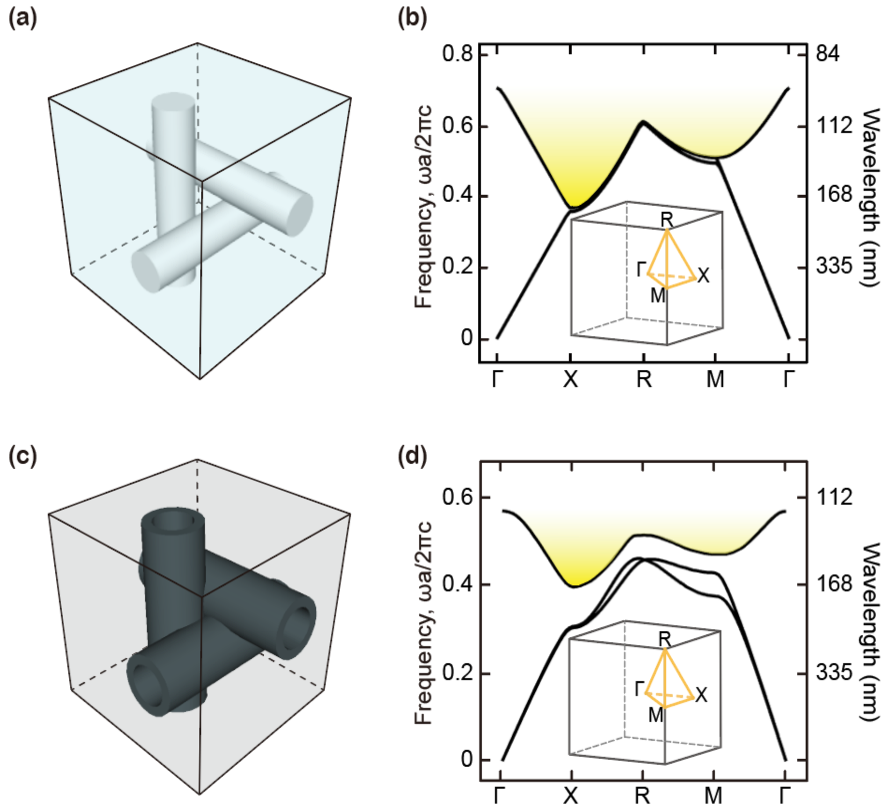


**Figure S2.** (a) Schematic representation for three-axis tensegrity in water medium, which was previously materialized.[5] The cylinder diameter, length, and *n* were set by 12.5 nm, 67nm, and 1.8, respectively. (b) Corresponding photonic band structure (DNA origami three-axis tensegrity in water medium). (c) Schematic representation for three-axis tensegrity with porous Si cladding in air medium, which could be developed after a post molding process. The cladding thickness and *n* were set by 10 nm and 3.0, respectively. (d) Corresponding photonic band structure. (b, d) All eigenfrequencies were non-dimensionalized with respect to the lattice constant ($a$ = 67 nm)

6. The pipeline for optimal bandgap engineering and corresponding results

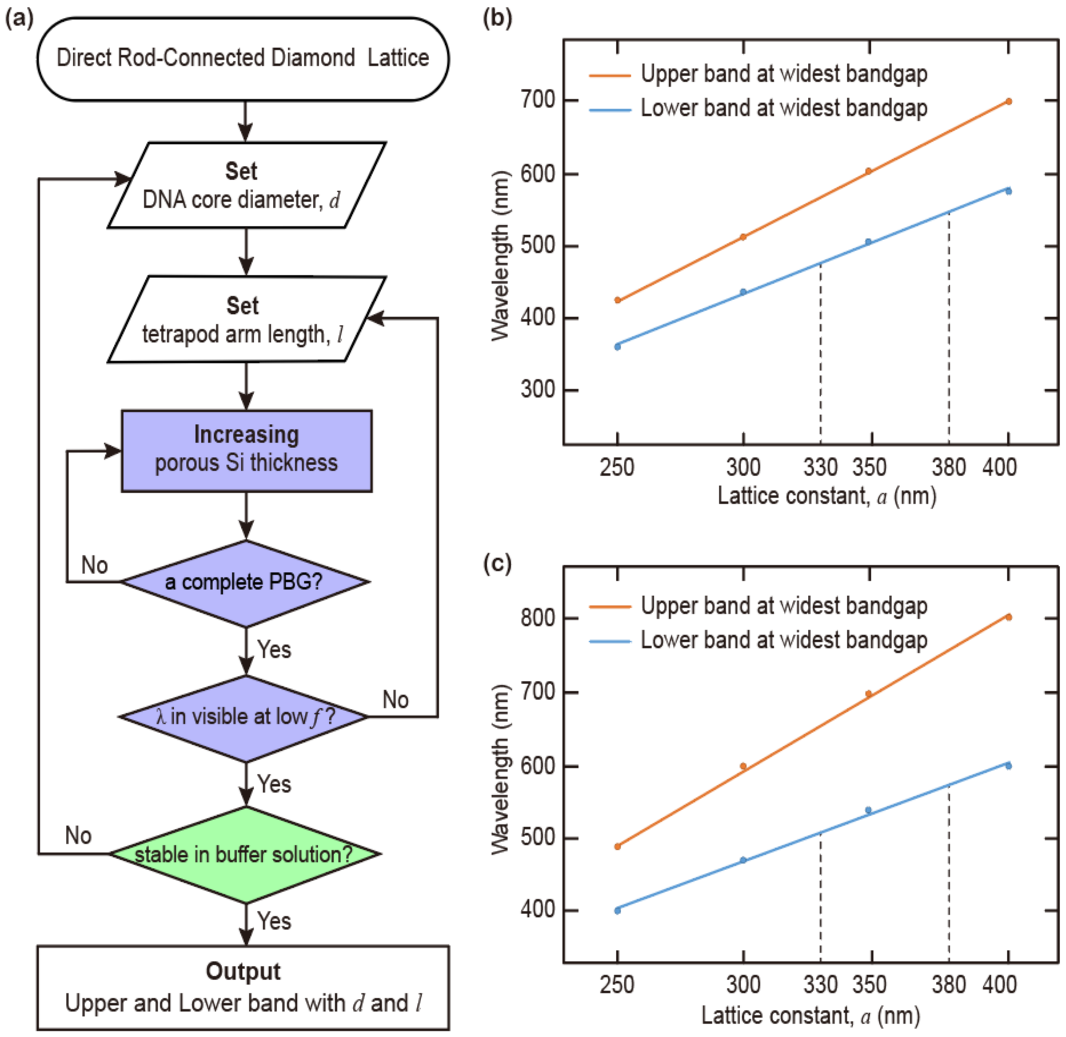


**Figure S3.** (a) The pipeline diagram for optimizing structural parameters. (b, c) The difference between the optimized upper and lower bands ($\delta$) according to the lattice constant ($a$). The cladding layer was assumed to be (b) porous silicon and (c) pure germanium.

### 7. Design diagram for simulation units

#### 7-1. Simulation unit 6/10-HB tetrapod

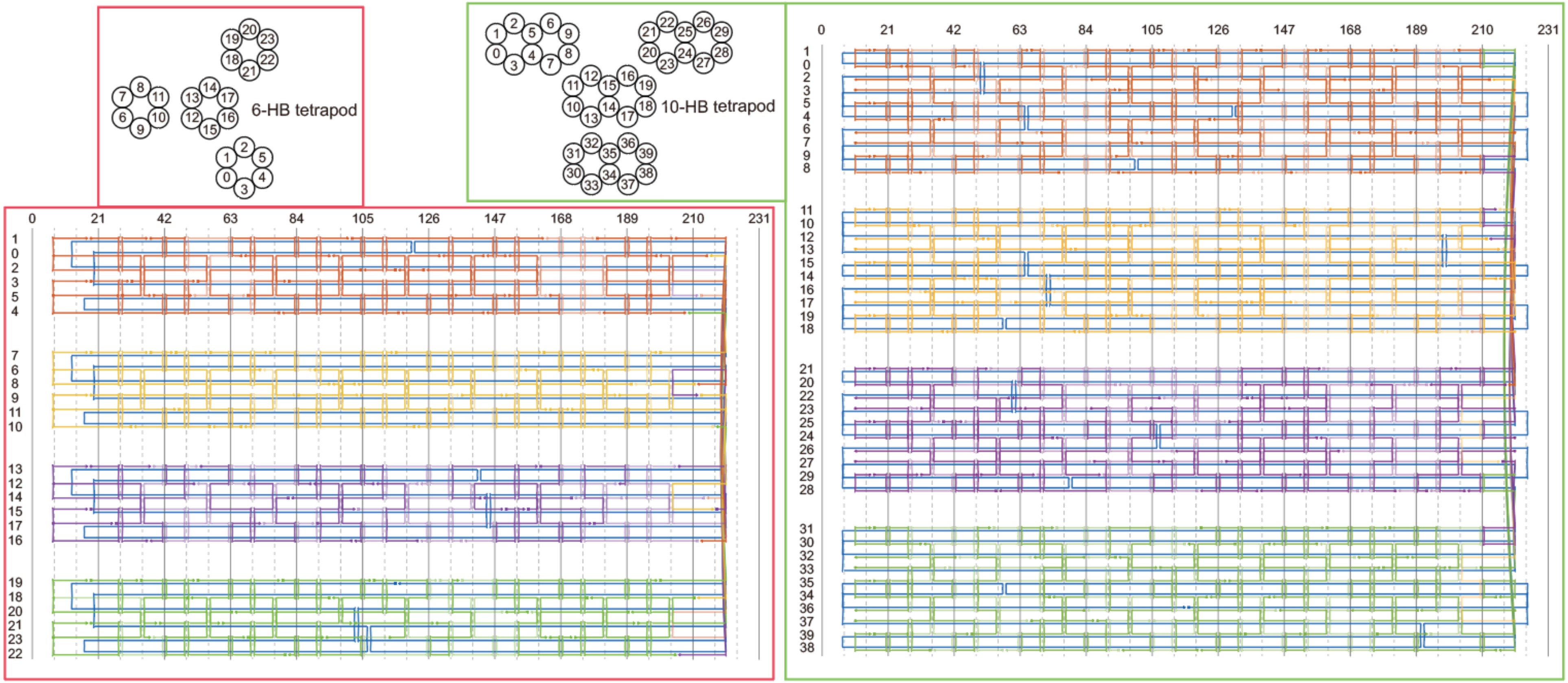


**Figure S4.** Detailed path of scaffolds (blue) and staples (other colors) using caDNAno.[2] The red box for 6-HB tetrapod cross-sectional view and design diagram. The green box for 10-HB tetrapod cross-sectional view and design diagram.

### 7-2. Simulation unit A

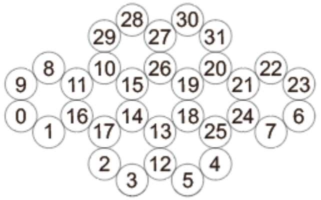


**Figure S5.** Detailed path of scaffolds (blue) and staples (gray and orange) using caDNAno.[2] The staples for gray represent shape-complementary binding site and the staples for orange represent bending site, which contribute to make 109.5°-angled unit, via insertions and deletions.

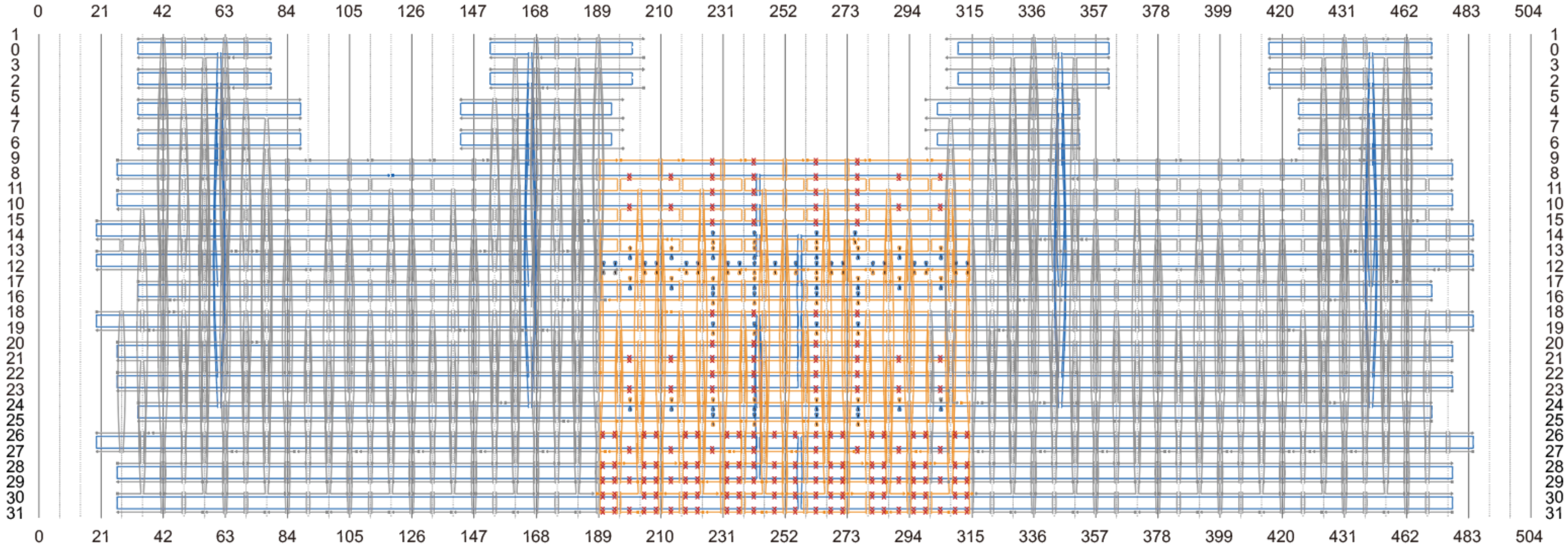

### 7-3. Simulation unit B

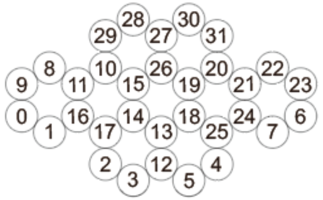


**Figure S6.** Detailed path of scaffolds (blue) and staples (gray and orange) using caDNAno.[2] The staples for gray represent shape-complementary binding site and the staples for orange represent bending site, which contribute to make 109.5°-angled unit, via insertions and deletions.

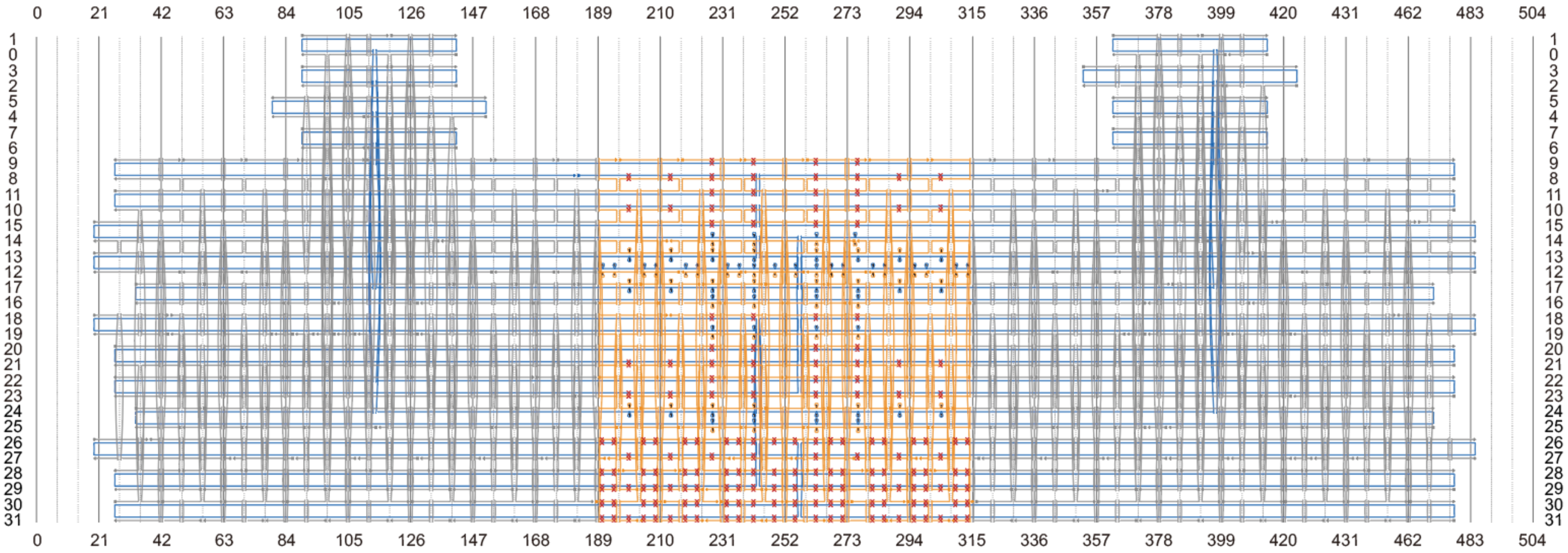

### 7-4. Simulation unit C

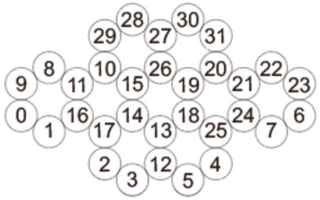


**Figure S7.** Detailed path of scaffolds (blue) and staples (gray and orange) using caDNAno.[2] The staples for gray represent shape-complementary binding site and the staples for orange represent bending site, which contribute to make 109.5°-angled unit, via insertions and deletions.

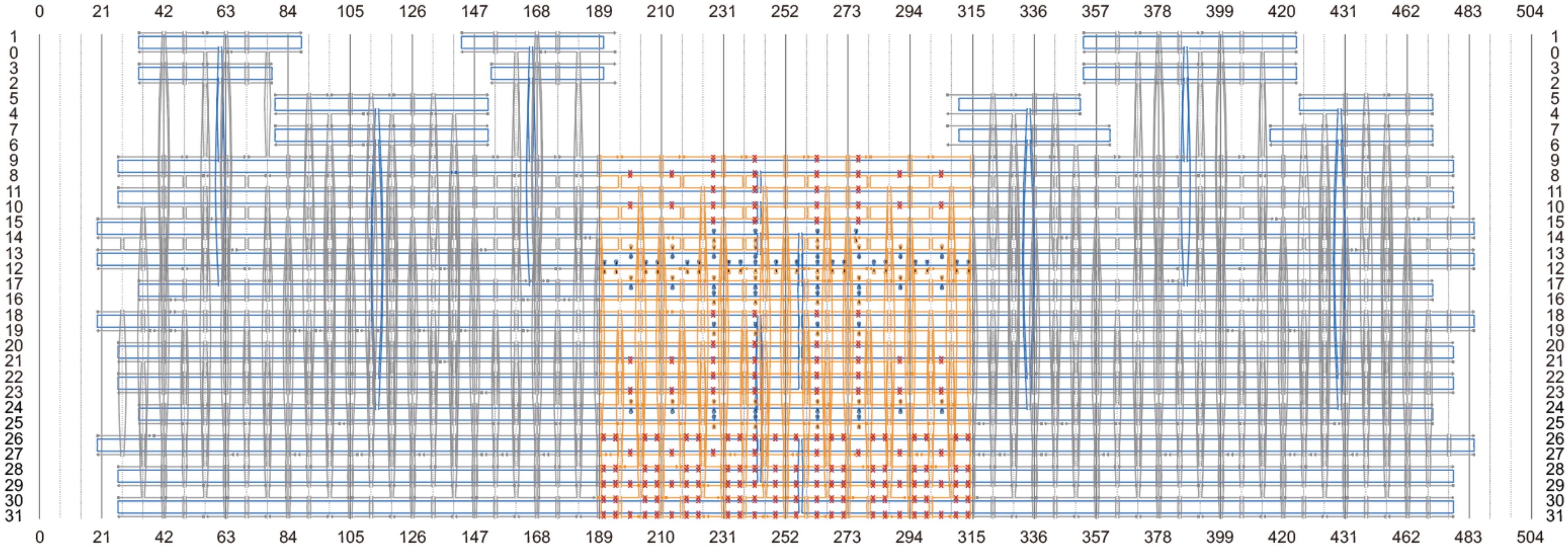

### Referneces


(1) Hur, K.; Hennig, R. G.; Wiesner, U., Exploring Periodic Bicontinuous Cubic Network Structures with Complete Phononic Bandgaps. *J. Phys. Chem. C* **2017**, *121*, 22347-22352.

(2) Douglas, S. M.; Marblestone, A. H.; Teerapittayanon, S.; Vazquez, A.; Church, G. M.; Shih, W. M., Rapid Prototyping of 3D DNA-Origami Shapes with caDNAno. *Nucleic Acids Res* **2009**, *37*, 5001-5006.

(3) Castro, C. E.; Kilchherr, F.; Kim, D.-N.; Shiao, E. L.; Wauer, T.; Wortmann, P.; Bathe, M.; Dietz, H., A Primer to Scaffolded DNA Origami. *Nat. Methods* **2011**, *8*, 221-229.

(4) Kim, D.-N.; Kilchherr, F.; Dietz, H.; Bathe, M., Quantitative Prediction of 3D Solution Shape and Flexibility of Nucleic Acid Nanostructures. *Nucleic Acids Res* **2011**, *40*, 2862-2868.

(5) Zhang, T.; Hartl, C.; Frank, K.; Heuer-Jungemann, A.; Fischer, S.; Nickels, P. C.; Nickel, B.; Liedl, T., 3D DNA Origami Crystals. *Adv. Mater.* **2018**, *30*, 1800273.
